# Fast and optimal sequence-to-graph alignment guided by seeds

**DOI:** 10.1101/2021.11.05.467453

**Authors:** Pesho Ivanov, Benjamin Bichsel, Martin Vechev

## Abstract

We present a novel A^⋆^ *seed heuristic* that enables fast and optimal sequence-to-graph alignment, guaranteed to minimize the edit distance of the alignment assuming non-negative edit costs.

We phrase optimal alignment as a shortest path problem and solve it by instantiating the A^⋆^ algorithm with our seed heuristic. The seed heuristic first extracts non-overlapping substrings (*seeds*) from the read, finds exact seed *matches* in the reference, marks preceding reference positions by *crumbs,* and uses the crumbs to direct the A^⋆^ search. The key idea is to punish paths for the absence of foreseeable seed matches. We prove admissibility of the seed heuristic, thus guaranteeing alignment optimality.

Our implementation extends the free and open source aligner and demonstrates that the seed heuristic outperforms all state-of-the-art optimal aligners including GraphAligner, Vargas, PaSGAL, and the prefix heuristic previously employed by AStarix. Specifically, we achieve a consistent speedup of >60× on both short Illumina reads and long HiFi reads (up to 25kbp), on both the *E. coli* linear reference genome (1Mbp) and the MHC variant graph (5Mbp). Our speedup is enabled by the seed heuristic consistently skipping >99.99% of the table cells that optimal aligners based on dynamic programming compute.

AStarix aligner and evaluations: https://github.com/eth-sri/astarix Full paper: https://www.biorxiv.org/content/10.1101/2021.11.05.467453

## 1 Introduction

Alignment of reads to a reference genome is an essential and early step in most bioinformatics pipelines. While linear references have been used traditionally, an increasing interest is directed towards graph references capable of representing biological variation [1]. Specifically, a *sequence-to-graph* alignment is a base-to-base correspondence between a given read and a walk in the graph. As sequencing errors and biological variation result in inexact read alignments, edit distance is the most common metric that alignment algorithms optimize in order to find the most probable read origin in the reference.

### Suboptimal alignment

In the last decades, approximate and alignment-free methods satisfied the demand for faster algorithms which process huge volumes of genetic data [2]. *Seed-and-extend* is arguably the most popular paradigm in read alignment [3, 4, 5]. First, substrings (called *seeds* or *kmers*) of the read are extracted, then aligned to the reference, and finally prospective matching locations are *extended* on both sides to align the full read.

While such a heuristic may produce acceptable alignments in many cases, it fundamentally does not provide quality guarantees, resulting in suboptimal alignment accuracy. In contrast, we demonstrate in this work that seeds can benefit optimal alignment as well.

### Key challenges in optimal alignment

Finding optimal alignments is desirable but expensive in the worst case, requiring 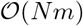 time [6], for graph size *N* and read length *m*. Unfortunately, most optimal sequence-to-graph aligners rely on dynamic programming (DP) and always reach this worst-case asymptotic runtime. Such aligners include Vargas [7], PaSGAL [8], GraphAligner [9], HGA [10], and VG [1], which use bit-level optimizations and parallelization to increase their throughput.

In contrast, AStarix [11] follows the promising direction of using a heuristic to avoid worst-case runtime on realistic data. To this end, AStarix rephrases the task of alignment as a shortest-path problem in an *alignment graph* extended by a *trie index,* and solves it using the A^⋆^ algorithm instantiated with a problem-specific prefix heuristic. Importantly, its choice of heuristic only affects performance, not optimality. Unlike DP-based algorithms, this prefix heuristic allows scaling sublinearly with the reference size, substantially increasing performance on large genomes. However, it can only efficiently align reads of limited length.

### This work: Optimal alignment for short and long reads

In this work, we address the key challenge of scaling to long HiFi reads, while retaining the superior scaling of AStarix in the size of the reference graph. To this end, we instantiate the A^⋆^ algorithm with a novel seed heuristic, which outperforms existing optimal aligners on both short and long HiFi reads. Specifically, the seed heuristic utilizes information from the whole read to narrowly direct the A^⋆^ search by placing *crumbs* on graph nodes which lead up to a *seed match,* i.e., an exact match of a substring of the read.

Overall, the contributions presented in this work are:

1. A novel A^⋆^ seed heuristic that exploits information from the whole read to quickly align it to a general graphs reference.
2. An optimality proof showing that the seed heuristic always finds an alignment with minimal edit distance.
3. An implementation of the seed heuristic as part of the AStarix aligner.
4. An extensive evaluation of our approach, showing that we align both short Illumina reads and long HiFi reads to both linear and graph references ≥ 60 × faster than existing optimal aligners.
5. A demonstration of superior empirical runtime scaling in the reference size *N*: *N*^0.46^ on Illumina reads and *N*^0.11^ on HiFi reads.

## 2 Prerequisites

We define the task of alignment as a shortest path problem (§2.1) to be solved using the A^⋆^ algorithm (§2.2).

### 2.1 Problem statement: Alignment as shortest path

In the following, we formalize the task of optimally aligning a read to a reference graph in terms of finding a shortest path in an *alignment graph.* Our discussion closely follows [11, §2] and is in line with [12].

#### Reference graph

A reference graph *G*_r_ = (*V*_r_, *E*_r_) encodes a collection of references to be considered when aligning a read. Its directed edges *E*_r_ ⊆ *V*_r_ × *V*_r_ × *Σ* are labeled by nucleotide letters from *Σ* = `{A, C, G, T}`, hence any walk *π*_r_ in *G*_r_ spells a string *σ*(*π*_r_) ∈ *Σ*\*.

An alignment of a read *q* ∈ *Σ*\* to a reference graph *G*_r_ consists of (i) a walk *π*_r_ in *G*_r_ and (ii) a sequence of edits (matches, substitutions, deletions, and insertions) that transform *σ*(*π*_r_) to *q*. An alignment is *optimal* if it minimizes the sum of edit costs for a given real-valued cost model *Δ* = (*Δ*_match_, *Δ*_subst_, *Δ*_del_, *Δ*_ins_). Throughout this work, we assume that edit costs are non-negative—a pre-requisite for the correctness of A^⋆^. Further, we assume that *Δ*_match_ ≤ *Δ*_subst_, *Δ*_ins_, *Δ*_del_—a prerequisite for the correctness of our heuristic.

We note that our approach naturally works for cyclic reference graphs.

#### Alignment graph

In order to formalize optimal alignment as a shortest path finding problem, we rely on an *alignment graph* 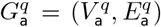. Its nodes 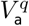 are *states* of the form 〈*v*, *i*〉, where *v* ∈ *V*_r_ is a node in the reference graph and *i* ∈ {0,…, |*q*|} corresponds to a position in the read *q*. Its edges 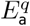 are selected such that any path *π*_a_ in 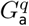 from 〈*u*, 0〉 to 〈*v*, *i*〉 corresponds to an alignment of the first *i* letters of *q* to *G*_r_. Further, the edges are weighted, which allows us to define an *optimal alignment* of a read *q* ∈ *Σ*\* as a shortest path *π*_a_ in 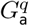 from 〈*u*, 0〉 to 〈*v*, |*q*|〉, for any *u*,*v* ∈ *V*_r_. Fig. 1 formally defines the edges 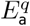.

**Fig. 1:**
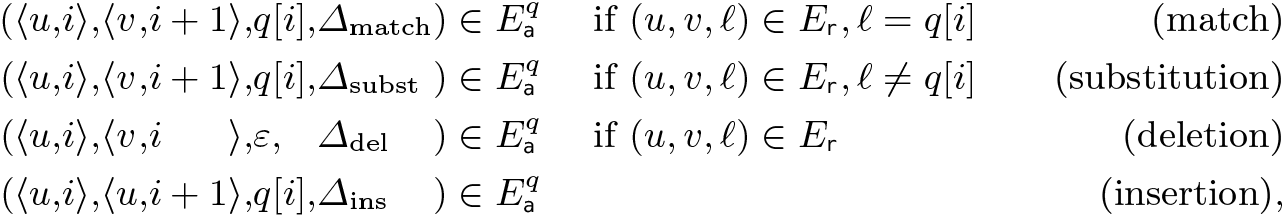
Formal definition of alignment graph edges 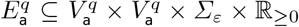. Here, *u*,*v* ∈ *V_r_*, 0 ≤ *i* < |*q*|, *ℓ* ∈ *Σ*, and *ε* represents the empty string, indicating that letter *ℓ* was deleted.

### 2.2 A^⋆^ algorithm for finding a shortest path

The A^⋆^ algorithm is a shortest path algorithm that generalizes Dijkstra’s algorithm by directing the search towards the target. Given a weighted graph *G* = (*V*, *E*), the A^⋆^ algorithm finds a shortest path from sources *S* ⊆ *V* to targets *T* ⊆ *V*. To prioritize paths that lead to a target, it relies on an admissible heuristic function 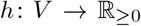, where *h*(*v*) estimates the remaining length of the shortest path from a given node *v* ∈ *V* to a target *t* ∈ *T*.

### Algorithm

In a nutshell, the A^⋆^ algorithm maintains a set of *explored* nodes, initialized by all possible starting nodes *S*. It then iteratively *expands* the explored state *v* with lowest estimated total cost *f* (*v*) by exploring all its neighbors. Here, *f*(*v*):= *g*(*v*) + *h*(*v*), where *g*(*v*) is the distance from *s* ∈ *S* to *v*, and *h*(*v*) is the estimated distance from *v* to *t* ∈ *T*. When the A^⋆^ algorithm expands a target node *t* ∈ *T*, it reconstructs the path leading to *t* and returns it. For completeness, App. A.1 provides an implementation of A^⋆^.

### Admissibility

The A^⋆^ algorithm is guaranteed to find a shortest path if its heuristic *h* provides a lower bound on the distance to the closest target, often referred to as *h* being *admissible* or optimistic.

Further, the performance of the A^⋆^ algorithm relies critically on the choice of *h*. Specifically, it is crucial to have low estimates for the optimal paths but also to have high estimates for suboptimal paths.

### Discussion

To summarize, we use the A^⋆^ algorithm to find a shortest path from 〈*u*, 0〉 to 〈*v*, |*q*|〉 in 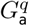. To guarantee optimality, its heuristic function *h*〈*v*, *i*〉 must provide a lower bound on the shortest distance from state 〈*v*, *i*〉 to a terminal state of the form 〈*w*, |*q*|〉. Equivalently, *h*〈*v*, *i*) should lower bound the minimal cost of aligning *q*[*i*:] to *G*_r_ starting from *v*, where *q*[*i*:] denotes the suffix of *q* starting at position *i* (0-indexed). The key challenge is thus finding a heuristic that is not only admissible but also yields favorable performance.

## 3 Seed heuristic

We instantiate the A^⋆^ algorithm with a novel, domain-specific *seed heuristic* which allows to quickly align reads to a general reference graph. We first intuitively explain the seed heuristic and showcase it on a simple example (§3.1). Then, we formally define the heuristic and prove its admissibility (§3.2). Finally, we adapt our approach to rely on a trie, which leads to a critical speedup (§3.3).

### 3.1 Overview

Fig. 2 showcases the seed heuristic on an overview example. It shows a read *q* to be aligned to a reference graph *G*_r_. Our goal is to find an optimal alignment starting from an arbitrary node *v* ∈ *G*_r_. For simplicity of the exposition, we assume unit edit costs *Δ* = (0,1,1,1), which we generalize in §3.2.

**Fig. 2:**
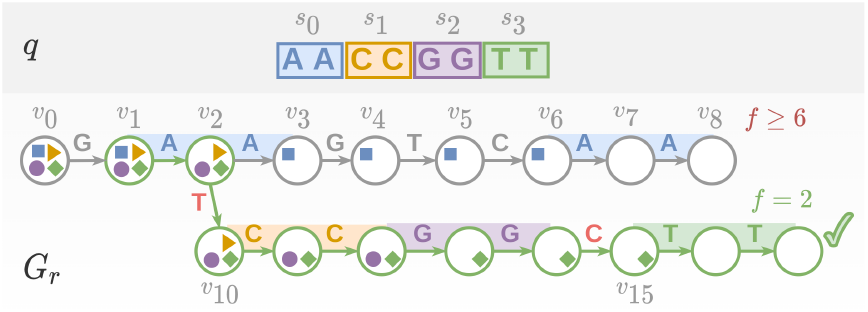
A toy overview example using the seed heuristic to align a read *q* to a reference graph *G_r_*. The read is split into four colored seeds, where their corresponding crumbs are shown inside reference graph nodes as symbols with matching color. The optimal alignment is highlighted as a green path ending with a tick 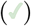 and includes one substitution (`T→A`) and one deletion (`C`).

#### Intuition

The A^⋆^ search requires us to provide a lower bound of the remaining path cost from a state 〈*v*, *i*〉 to a target state. Clearly, to align the whole query, each of the remaining seeds (i.e. at or after position *i* in *q*) has to be eventually aligned. The intuition underlying the seed heuristic is to punish the state for the absence of any foreseeable match of each remaining seed. Notice that the order of the seeds is not directly taken into account.

In order to quickly check if a seed *s* can lead to a match, we follow a procedure similar to the one used by Hansel and Gretel who were placing breadcrumbs to find their trail back home. Before aligning a query, we will precompute all *crumbs* from all seeds so that not finding a crumb for a seed *s* on node *v* indicates that seed *s* could not be matched exactly before the query is fully aligned continuing from *v*. This way, assuming that a shortest path includes many seed matches, the crumbs will direct the A^⋆^ search along with it.

If a crumb from an expected seed is missing in node *v*, its corresponding seed *s* could not possibly be aligned exactly and this will incur the cost of at least one substitution, insertion, or deletion. Assuming unit edit costs, *h*〈*v*, *i*〉 yields a lower bound on the cost for aligning *q*[*i*:] starting from *v* by simply returning the number of missing expected crumbs in *v*.

#### Crumbs precomputation example

Fig. 2 shows four seeds as colored sections of length 2 each, and represents their corresponding crumbs as 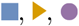 and 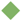, respectively. Four crumbs are expected if we start at *v*_2_, but 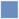 is missing, so *h*〈*v*_2_, 0〉 = 1. Analogously, if we reach *v*_2_ after aligning one letter from the read, we expect 3 crumbs (except 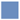), and we find them all in *v*_2_, so *h*〈*v*_2_, 1〉 =0. To precompute the crumbs for each seed, we first find all positions in *G*_r_ from which the seed aligns exactly. Fig. 2 shows these exact matches as colored sections of *G*_r_. Then, from each match we traverse *G*_r_ backwards and add crumbs to nodes that could lead to these matches. For example, because seed `CC` can be matched starting in node *v*_10_, crumbs 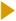 are placed on all nodes leading up to *v*_10_. Similarly, seed `AA` has two exact matches, one starting in node *v*_0_ and one starting in node *v*_6_. However, we only add crumbs 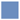 to nodes *v*_0_, *v*_1_, and *v*_3_–*v*_6_, but not to node *v*_2_. This is because *v*_2_ is (i) strictly after the beginning of the match of `AA` at *v*_1_ and (ii) too far before the match of `AA` at v_6_. Specifically, any alignment starting from node *v*_2_ and still matching `AA` at *v*_6_ would induce an overall cost of 4 (it would require deleting the 4 letters *A*, *G*, *T*, and *C*). Even without a crumb 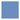 on *v*_2_, our heuristic provides a lower bound on the cost of such an alignment: it never estimates a cost of more than 4, the number of seeds.

#### Guiding the search example

Fig. 3 demonstrates how *h*〈*v*, *i*〉 guides the A^⋆^ algorithm towards the shortest path by showing which states may be expanded when using the seed heuristic. Specifically, the unique optimal alignment in Fig. 2 starts from node *v*_1_, continues to *v*_2_, and then proceeds through node *v*_10_ (instead of *v*_3_).

**Fig. 3:**
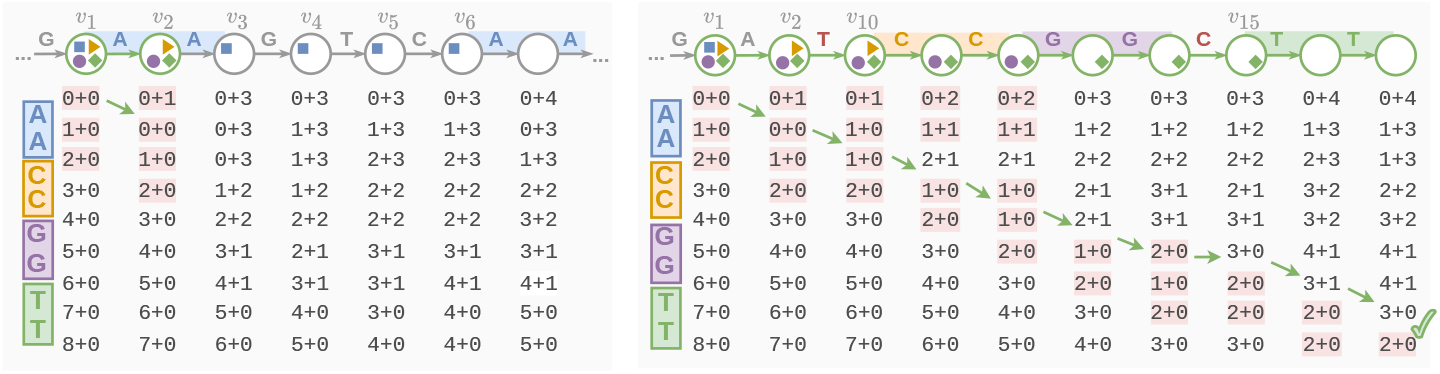
Exploration of 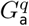, searching for a shortest path from the first to the last row using the seed heuristic. The table entry in the *i*^th^ row (zero indexed) below node *v* shows *g*〈*v*, *i*〉 + *h*〈*v*, *i*〉, where *g*〈*v*, *i*〉 is the shortest distance from any starting state 〈*u*, 0〉 to 〈*v*, *i*〉. States that may^1^ be expanded by the A^⋆^ algorithm are highlighted in pink, and the rest of the states are shown for completeness even though they are never expanded. The shortest path corresponding to the best alignment is shown with green arrows 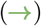.

While the seed heuristic initially explores all states of the form 〈*v*, 0〉 (we discuss in §3.3 how to avoid this by using a trie), it skips expanding any state that involves nodes *v*_3_–*v*_8_. This improvement is possible because all these explored states are penalized by the seed heuristic by at least 3, while the shortest path of cost 2 will be found before considering states on nodes *v*_3_–*v*_8_. Here, the heuristic function accurately predicts that expanding *v*_10_ may eventually lead to an exact alignment of seeds `CC`, `GG` and `TT`, while expanding *v*_3_ may not lead to an alignment of either seed. In particular, the seed heuristic is not misled by the short-term benefit of correctly matching *A* in *v*_2_, and instead provides a longterm recommendation based on the whole read. Thus, even though the walk to *v*_3_ aligns exactly the first two letters of *q*, A^⋆^ does not expand *v*_3_ because the seed heuristic guarantees that the future cost will be at least 3.

### 3.2 Formal definition

Next, we formally define the seed heuristic function *h*〈*v*, *i*〉. Overall, we want to ensure that *h*〈*v*, *i*〉 is admissible, i.e., that it is a lower bound on the cost of a shortest path from 〈*v*, *i*〉 to some 〈*w*, |*q*|〉 in 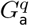.

#### Seeds

We split read *q* ∈ *Σ*\* into a set *Seeds* of non-overlapping seeds *s*_0_,…, *s*_|*Seeds*|–1_ ∈ Σ*. For simplicity, in this work we ensure that all seeds have the same length and are consecutive, i.e., we split *q* into substrings *s*_0_ · *s*_1_ · · · *s*_|*Seeds*|–1_ · *t*, where all *s_j_* are seeds of length *k* and we ignore the suffix *t* of *q*, which is shorter than *k*. We note that our approach could be trivially generalized to seeds of different lengths or non-consecutive seeds as long as they do not overlap. An interesting future work item is investigating how different choices of seeds affect the performance of our approach, and selecting seeds accordingly.

#### Matches

For each seed *s* ∈ *Seeds*, we locate all nodes *u* ∈ *M*(*s*) in the reference graph that can be the start of an exact match of *s*:

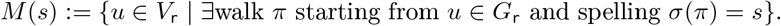

To compute *M*(*s*) efficiently, we leverage the trie introduced in §3.3.

#### Crumbs

For seed *s_j_* starting at position *i* in *q*, we place crumbs on all nodes *u* ∈ *V*_r_ which can reach a node *v* ∈ *M*(*s_j_*) using less than *i* + *n*_del_ edges:

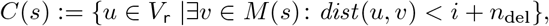

where *dist*(*u*, *v*) is the length of a shortest walk from *u* to *v*.

Later in this section, we will select *n*_del_ to ensure that if an alignment uses more than *n*_del_ deletions, its cost must be so high that the heuristic function is trivially admissible.

To compute *C*(*s*) efficiently, we can traverse the reference graph backwards from each *v* ∈ *M*(*s*) by a backward breadth-first-search (BFS).

#### Heuristic

Let *Seeds*_≥*i*_ be the set of seeds that start at or after position *i* of the read, formally defined by *Seeds*_≥*i*_:= {*s_j_* | ⌈*i*/*k*⌉ ≤ *j* 〈 |*Seeds*|}. This allows us to define the number of expected but missing crumbs in state 〈*v*, *i*〉 as *misses*〈*v*, *i*〉:= |{*v* ∉ *C*(*s*) | *s* ∈ *Seeds*_≥*i*_}|. Finally, we define the seed heuristic as

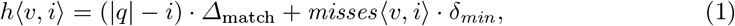

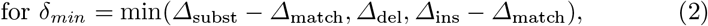

Intuitively, Eq. (1) reflects that the cost of aligning each remaining letter from *q*[*i*:] is at least *Δ*_match_. In addition, every inexact alignment of a seed induces an additional cost of at least *δ_min_*. Specifically, every substitution costs *Δ*_subst_ but requires one less match; every deletion costs *Δ*_del_; and every insertion costs *Δ*_ins_ but also requires one less match.

We note that *h*〈*v*, *i*〉 implicitly also depends on the reference graph *G*_r_, the read *q*, the set of seeds, and the edit costs *Δ*.

In order for an alignment with at least *n_del_* deletions to have a cost so high that the heuristic function is trivially admissible, we ensure *n_del_* · *Δ_del_* ≥ *h*〈*v*, *i*〉 by defining

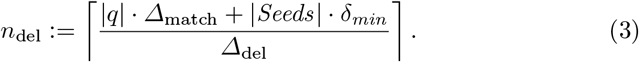

In Thm. 1, we show that *h*〈*v*, *i*〉 is admissible, ensuring that our heuristic yields optimal alignments.

##### Theorem 1 (Admissibility).

*The seed heuristic* h(v,i) *is admissible.*

*Proof.* We provide a proof for Thm. 1 in App. A.2.

### 3.3 Trie index

Considering all nodes *v* ∈ *V*_r_ as possible starting points for the alignment means that the A^⋆^ algorithm would explore all states of the form 〈*v*, 0〉, which immediately induces a high overhead of |*V*_r_|. In line with previous works [11, 13], we avoid this overhead by complementing the reference graph with a trie index to produce a new graph 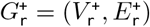, where 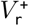 is the union of the reference graph nodes *V*_r_ and the new trie vertices, and 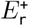 is the union of *E*_r_, the trie edges, and edges connecting the trie leafs with reference nodes. Note that constructing this trie index is a one-time pre-processing step that can be reused for multiple queries.

Since we want to also support aligning reverse-complement reads by starting from the trie *root*, we build the trie not only from the original reference graph and also from its reverse-complement.

#### Intuition

Fig. 4 extends the reference graph *G*_r_ from Fig. 2 with a trie. Here, any path in the reference graph uniquely corresponds to a path starting from the trie *root* (the top-most node in Fig. 4). Thus, in order to find an optimal alignment, it suffices to consider paths starting from the trie *root*, by using state *(root*, 0) as the only source for the A^⋆^ algorithm. Note that if the reference graph branches frequently, the number of paths with length D may rise exponentially, leading to an exponential number of trie leaves. To counteract this exponential growth, we can select *D* logarithmically small, as log_4_ *N*.

**Fig. 4:**
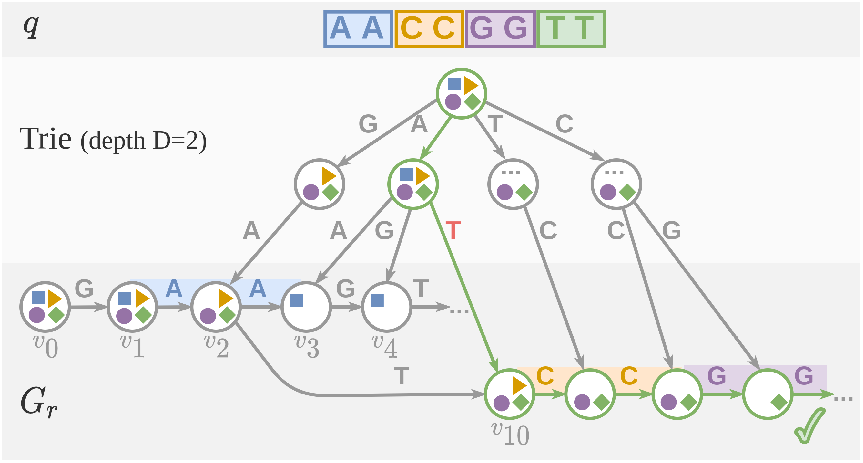
Reference graph from Fig. 2, extended by a trie of depth *D* = 2. For simplicity, the reverse-complement reference graph and parts marked by “…” are omitted.

For a more thorough introduction to the trie and its construction, see [11]. Importantly, our placement of crumbs (§3.2) generalizes directly to reference graphs extended with a trie (see also Fig. 4).

#### Reusing the trie to find seed matches

As a second usage of the trie, we can also exploit it to efficiently locate all matches *M*(*s*) of a given seed *s*. In order to find all nodes where a seed match begins, we align (without errors) 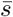, the reverse-complement of *s*. To this end, we follow all paths spelling 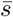 starting from the *root*—the final nodes of these paths then correspond to nodes in *M*(*s*). We ensure that the seed length |*s*| is not shorter than the trie depth *D*, so that matching all letters in 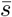 ensures that we eventually transition from a trie leaf to the reference graph.

#### Optimization: skip crumbs on the trie

Generally, we aim to place as few crumbs as possible, in order to both reduce precomputation time and avoid misleading the A^⋆^ algorithm by unnecessary crumbs. In the following, we introduce an optimization to avoid placing crumbs on trie nodes that are “too close” to the match of their corresponding seed so they cannot lead to an optimal alignment.

Specifically, when traversing the reference graph backwards to place crumbs for a match of seed s starting at node *w*, we may “climb” from a reference graph node u to a trie node *u*′ backwards through an edge that otherwise leads from the trie to the reference. Assuming s starts at position *i* in the read, we have already established that we can only consider nodes u that can reach *w* with less than *i* + *n*_del_ edges (see §3.2). Here, we observe that it is sufficient to only climb into the trie from nodes *u* that can reach *w* using more than *i* – *n*_ins_ – *D* edges, for

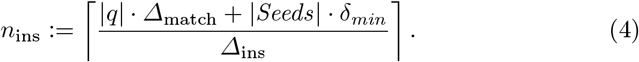

We define *n*_ins_ analogously to *n*_del_ to ensure that *n*_ins_ insertions will induce a cost that is always higher than *h*〈*u*, *i*〉. We note that we can only avoid climbing into the trie if all paths from *u* to *w* are too short, in particular the longest one.

The following Lemma 1 shows that this optimization preserves optimality.

##### Lemma 1 (Admissibility when skipping crumbs)

*The seed heuristic remains admissible when crumbs are skipped in the trie.*

*Proof.* We provide a proof for Lemma 1 in App. A.2.

In order to efficiently identify all nodes *u* that can reach *w* by using more than *i* — *D* — *n*_ins_ edges (among all nodes at a backward-distance at most *i* + *n*_del_ from *w*), we use topological sorting: considering only nodes at a backward-distance at most *i* + *n*_del_ from *w*, the length of a longest path from a node *v* to *w* is (i) ∞ if *v* lies on a cycle and (ii) computable from nodes closer to *w* otherwise.

## 4 Evaluations

In the following, we demonstrate that our approach aligns faster than existing optimal aligners due to its superior scaling. Specifically, we address the following research questions:

**Q1** What speedup can the seed heuristic achieve?
**Q2** How does the seed heuristic scale with reference size?
**Q3** How does the seed heuristic scale with read length?

The modes of operation which we analyze include both short (Illumina) and long (HiFi) reads to be aligned on on both linear and graph references.

### 4.1 Seed heuristic implementation

Both the seed heuristic and the prefix heuristic reuse the same free and open source C++codebase of the AStarix aligner [11]. It includes a simple implementation of a graph and trie data structure which is not optimized for memory usage. In order to easily align reverse complement reads, the reverse complement of the graph is stored alongside its straight version. The shortest path algorithm only constructs explored states explicitly, so most states remain implicitly defined and do not cause computational burden.

Both heuristics benefit from a default optimization in AStarix called *greedy matching* [11, Section 4.2] which skips adding a state to the A^⋆^ queue when only one edge is outgoing from a state and the upcoming read and reference letters match.

### 4.2 Setting

All experiments were executed on a commodity machine with an Intel Core i7-6700 CPU @ 3.40GHz processor, using a memory limit of 20 GB and a single thread. We note that while multiple tools support parallelization when aligning a single read, all tools can be trivially parallelized to align multiple reads in parallel.

#### Compared aligners

We compare the novel seed heuristic to prefix heuristic (both heuristics are implemented in AStarix), GraphAligner, PaSGAL, and Vargas. We provide the versions and commands used to run all aligners and read simulators in App. A.3. We note that we do not compare to VG [1] and HGA [10] since the optimal alignment functionality of **VG** is meant for debugging purposes and has been shown to be inferior to other aligners [10, Tab. 4], and **HGA**makes use of parallelization and a GPU but has been shown to be superseded in the single CPU regime [10, Fig. 9]. PaSGAL and Vargas are compiled with AVX2 support. We execute the prefix heuristic with the default lookup depth *d* = 5.

#### Seed heuristic parameters

The choice of parameters for the seed heuristic influences its performance. Increasing the trie depth increases its number of nodes, but decreases the average out-degree of its leaves. We set the trie depth for all experiments to *D* = 14 ≈ log_4_ *N*.

Shorter seeds are more likely to be matched perfectly by an optimal alignment, as they contain less letters that could be subject to edits. Thus, shorter seeds can tolerate higher error rate, but at the cost of slower precomputation due to a higher total number of matches, and slower alignment due to more off-track matches. In our experiments, we use seed lengths of *k* = 25 for Illumina reads and *k* = 150 for HiFi reads.

#### Data

We aligned reads to two different reference graphs: a linear *E. coli* genome (strain: K-12 substr. MG1655, ASM584v2) [14], with length of 4641652bp (approx. 4.7Mbp), and a variant graph with the Major Histocompatibility Complex (MHC), of size 5 318019bp (approx. 5Mbp), taken from the evaluations of PaSGAL [8]. Additionally, we extracted a path from MHC in order to create a linear reference MHC-linear of length 4 956 648bp which covers approx. 93% of the original graph. Because of input graph format restrictions, we execute GraphAligner, Vargas and PaSGAL only on linear references in FASTA format *(E. coli* and the MHC-linear), while we execute the seed heuristic and the prefix heuristic on the original references *(E. coli* and MHC). This yields an underestimation of the speedup of the seed heuristic, as we expect the performance on MHC-linear to be strictly better than on the whole MHC graph.

To generate both short Illumina and long HiFi reads, we relied on two tools. We generated short single-end 200bp Illumina MSv3 reads using **ART** simulator [15]. We generated long HiFi reads using the script randomreads.sh^2^ with sequencing lengths 5-25kbp and error rates 0.3%, which are typical for HiFi reads.

#### Edit costs

We execute AStarix with edit costs typical to the corresponding sequencing technology: *Δ* = (0,1, 5, 5) for Illumina reads and *Δ* = (0,1,1,1) for HiFi reads. As the performance of DP-based tools is independent of edit costs, we are using the respective default edits costs when executing GraphAligner, PaSGAL and Vargas.

#### Metrics

We report the performance of aligners in terms of runtime per one thousand aligned base pairs [s/kbp]. Since we measured runtime end-to-end (in-cluding loading reference graphs and reads from disk, and building the trie index for AStarix), we ensured that alignment time dominates the total runtime by providing sufficiently many reads to each aligner. In order to prevent excessive runtimes for slower tools, we used a different number of reads for each tool and explicitly report them for each experiment.

Since shortest path approaches skip considerable parts of the computation performed by aligners based on dynamic programming, the commonly used Giga Cell Updates Per Second (GCUPS) metric is not adequate for measuring performance in our context.

We measured used memory by `max_rss` (Maximum Resident Set Size) from Snakemake^3^.

We do not report accuracy or number of unaligned reads, as all evaluated tools align all reads with guaranteed optimality according to edit distance. We note that Vargas reports a warning that some of its alignments are not optimal—we ignore this warning and focus on its performance.

### 4.3 Q1: Speedup of the seed heuristic

Table 1 shows that the seed heuristic achieves a speedup of at least 60 times compared to all considered aligners, across all regimes of operation: both Illumina and HiFi reads aligned on *E. coli* and MHC references.

**Table 1:**
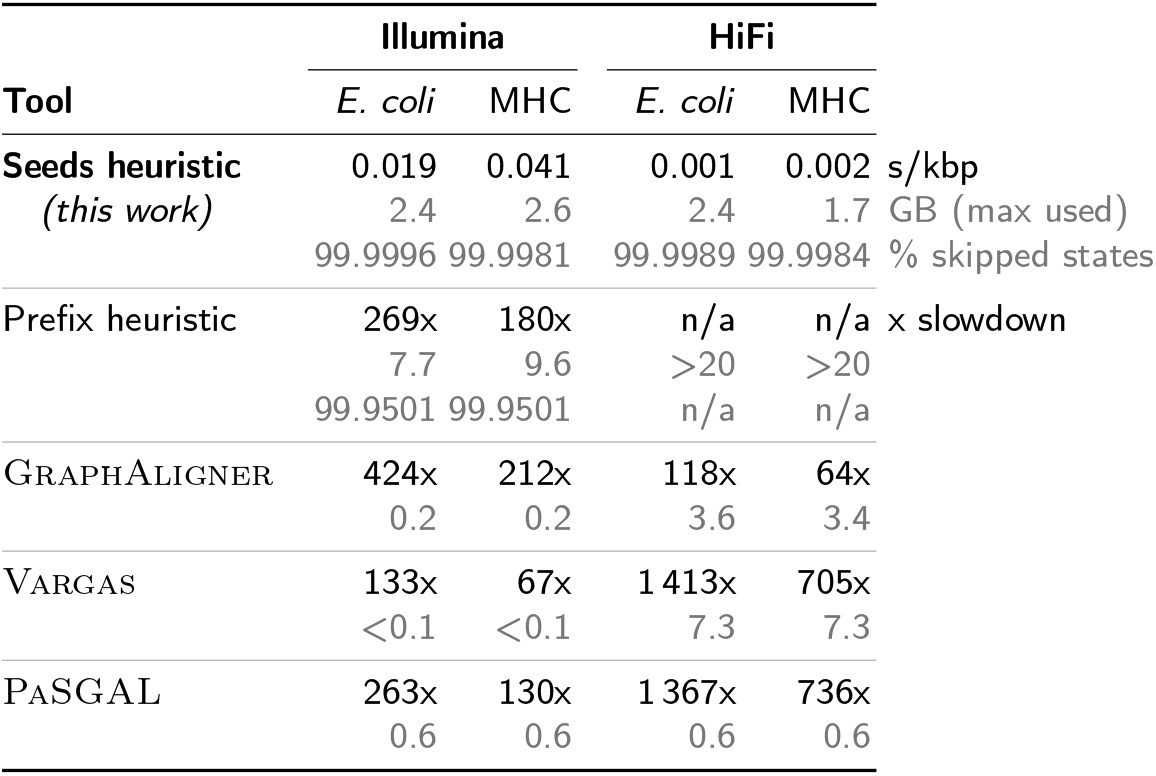
Runtime and memory comparison of optimal aligners. Simulated Illumina and HiFi reads are aligned to linear *E. coli* and graph MHC references. The runtime of the seed heuristic is expressed as absolute time per aligned kbp, while the other aligners are compared to the seed heuristic at a fold change. Additionally, the fraction of explored states is shown for the seed heuristic and the prefix heuristic.

In the Illumina experiments, the seed heuristic is given 100k reads, while the other tools are given 1000 reads. In the HiFi experiments, the seed heuristic is given reads that cover the reference 10 times, and the other tools are given reads of coverage 0.1.

The key reason for the speedup of the seed heuristic is that on all four experiments, it skips ≥ 99.99% of the *Nm* states computed by the DP approaches of GraphAligner, PaSGAL, and Vargas. This fraction accounts for both the explored states during the A^⋆^ algorithm, and the number of crumbs added to nodes during precomputation for each read.

The prefix heuristic exceeded the available memory on HiFi reads, as it is not designed for long reads.

### 4.4 Q2: Scaling with reference size

In order to study the scaling of the aligners in terms of the reference size, we extracted prefixes of increasing length from MHC-linear. We then generated reads from each prefix, and ran all tools on all prefixes with the corresponding reads.

#### Illumina reads

Fig. 5 shows the runtime scaling and memory usage for Illumina reads. The seed heuristic was provided with 10k reads, while other tools were provided with 1k reads. The runtime of GraphAligner, PaSGAL and Vargas grow approximately linearly with the reference length, whereas the runtime of the seed heuristic grows roughly with 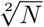, where *N* is the reference size. Even on relatively small graphs like MHC, the speedup of the seed heuristic reaches 200 times. Note that the scaling of the prefix heuristic is substantially worse than the seed heuristic since the 200bp reads are outside of its operational capabilities.

**Fig. 5:**
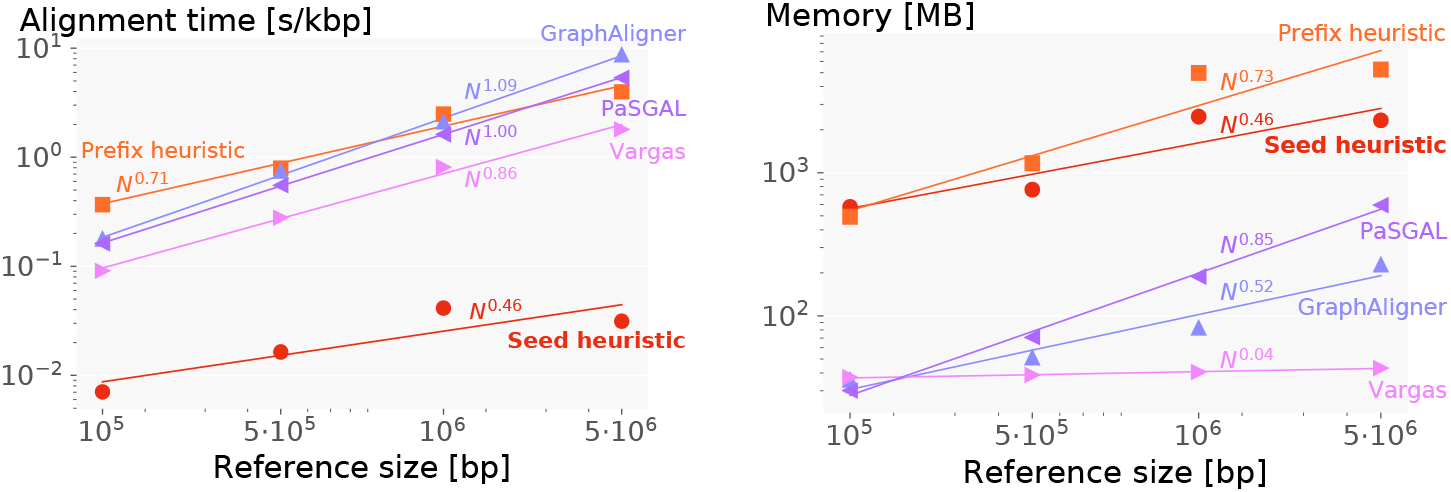
Performance degradation with reference size for **Illumina reads**. Log-log plots of total alignment time (left) and memory usage (right) show the scaling difference between aligners.

#### HiFi reads

Fig. 6 shows the runtime scaling and memory usage for HiFi reads. The respective total lengths of all aligned reads are 5Mbp for the seed heuristic, 500kbp for GraphAligner, and 100kbp for Vargas and PaSGAL. We do not show the prefix heuristic, since it explores too many states and runs out of memory. Crucially, we observe that the runtime of the seed heuristic is almost independent of the reference size, growing as *N*^0.11^. We believe this improved trend compared to short reads is because the seed heuristic obtains better guidance on long reads, as it can leverage information from the whole read.

**Fig. 6:**
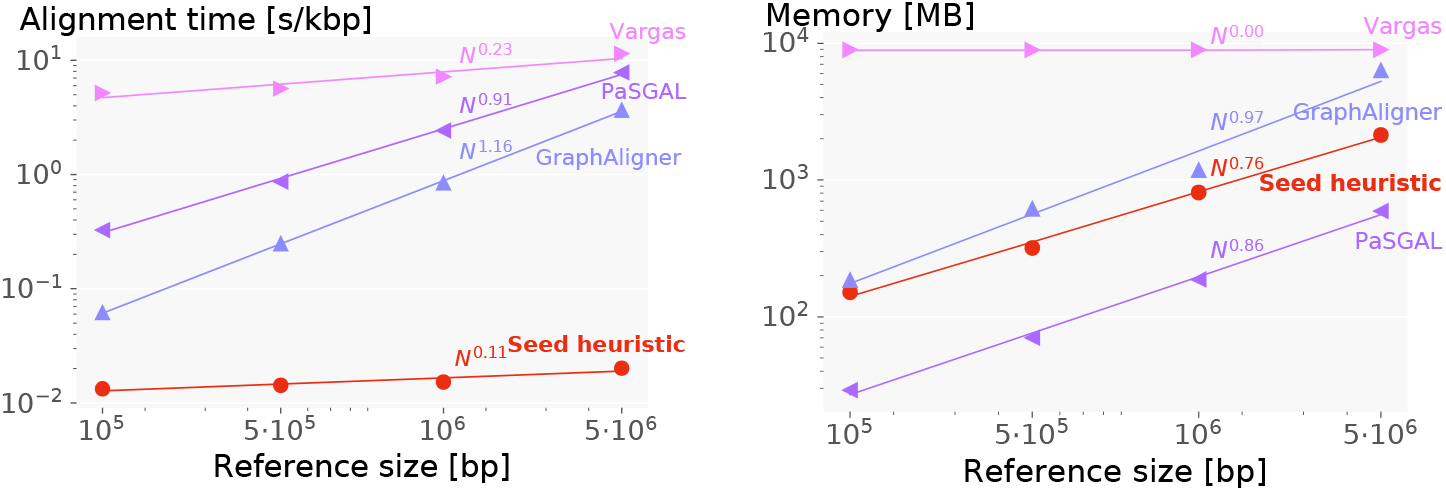
Performance degradation with reference size for **HiFi reads**. Log-log plots of total alignment time (left) and memory usage (right) show the scaling difference between aligners. Linear best fits correspond to polynomials of varying degree.

For both Illumina and HiFi reads, we observe near-linear scaling for PaSGAL and GraphAligner as expected from the theoretical 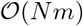 runtime of the DP approaches. We conjecture that the runtime of Vargas for long reads is dominated by the dependence from the read length, which is why on HiFi reads we observe better than linear runtime dependency on *N* but very large runtime. The current alignment bottleneck of AStarix-seeds is its memory usage, which is distributed between remembering crumbs and holding a queue of explored states.

### 4.5 Q3: Scaling with read length

Fig. 7 shows the runtime and memory scaling with increasing length of aligned HiFi reads on MHC reference. Here we used reads with a total length of 100Mbp for the seed heuristic and 2Mbp for all other aligners.

**Fig. 7:**
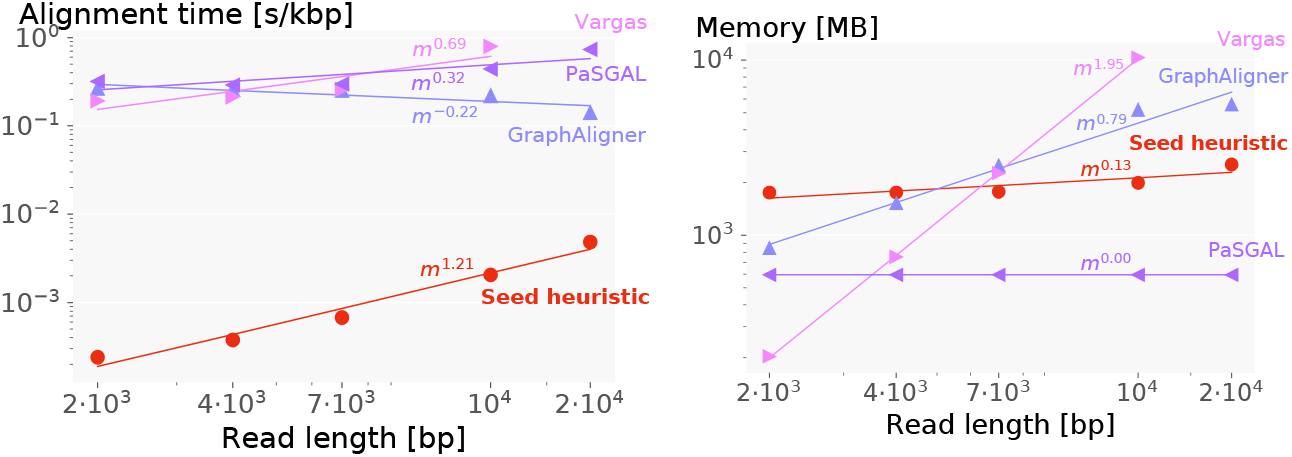
Performance degradation with HiFi read length. Log-log plots of total alignment time (left) and memory usage (right) show the scaling difference between aligners.

The scaling of the seed heuristic in terms of read length is slightly worse than that of other aligners. However, this is compensated by its superior scaling in terms of reference size (see §4.4), leading to an overall better absolute runtime.

We note that the memory usage of the seed heuristic does not heavily depend on the read length and for reads longer than 10kbp, it is superior to GraphAligner and Vargas.

## 5 Acknowledgements

We want to thank the anonymous reviewers for the valuable feedback. The first author is grateful for the Vipassana meditation which inspired the idea to punish paths for the absence of foreseeable seed matches.

## 6 Conclusion

We have presented an optimal read aligner based on the A^⋆^ algorithm instantiated with a novel seed heuristic which guides the search by preferring crumbs on nodes that lead towards optimal alignments even for long reads.

The memory usage is currently limiting the application of AStarix for bigger references due to the size of the trie index. A remaining challenge is designing a heuristic function able to handle not only long but also noisier reads, such as the uncorrected PacBio reads that may reach 20% of mistakes. Possible improvements of the seed heuristic may include inexact matching of seeds, careful choice of seed positions, and accounting for the seeds order.

## A Appendix

### Algorithm 1 A^⋆^ algorithm

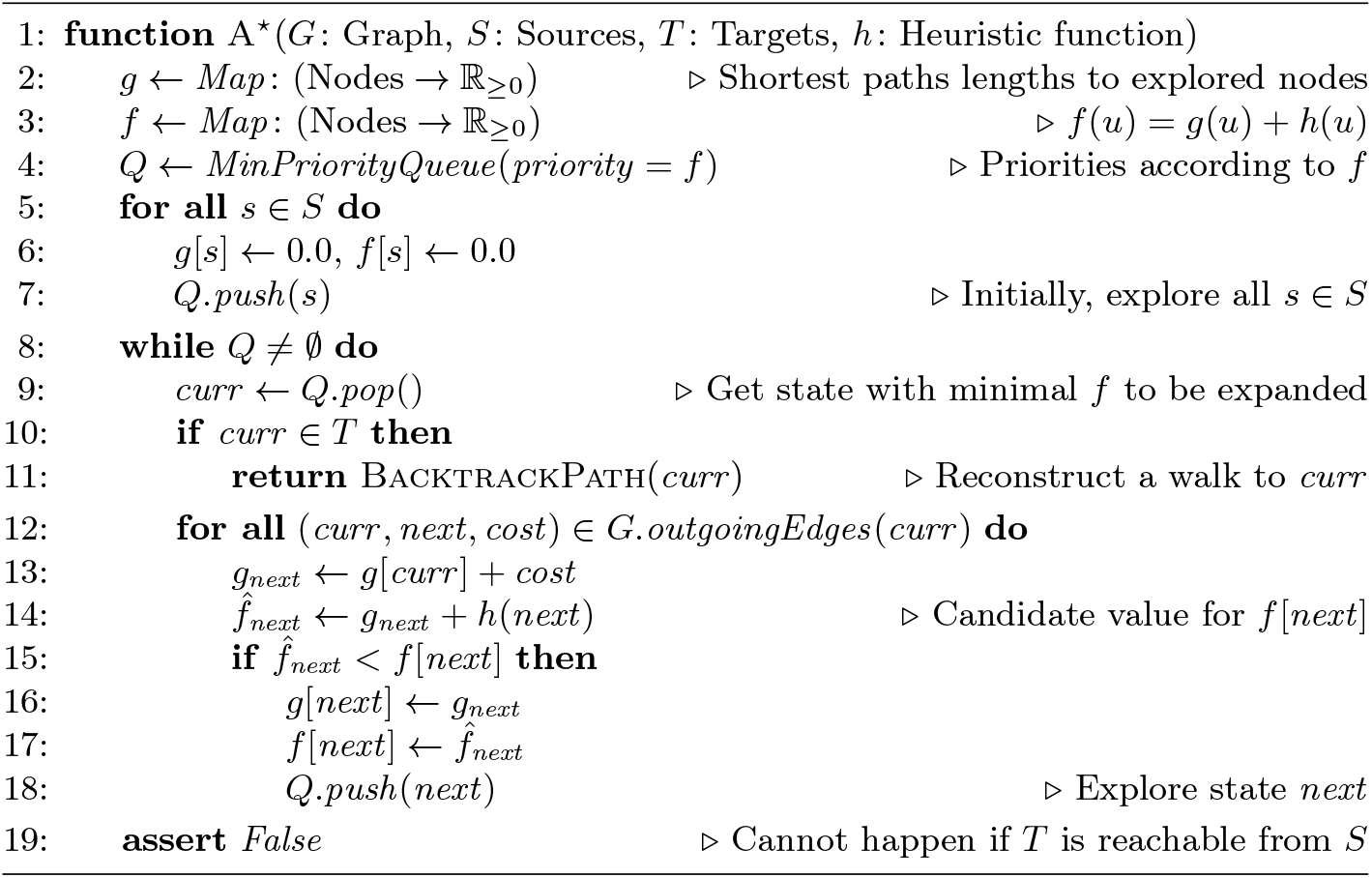

### A.1 A^⋆^ Algorithm

Algorithm 1 shows an implementation of the A^⋆^ algorithm, taken from [11, §A.1]. We omit the implementation of BacktrackPath for simplicity.

### A.2 Proofs

In the following, we provide proofs for Thm. 1 and Lemma 1, restated here for convenience.

#### Theorem 1 (Admissibility).

*The seed heuristic h*〈*v*,*i*〉 *is admissible.*

*Proof.* Let *A* be an optimal alignment of *q*[*i*:] starting from *v* ∈ *G*_r_. We will prove that the cost of *A* is at least *h*〈*v*, *i*〉.

If *A* contains at least *n*_del_ deletions, its cost is at least *n*_del_ · *Δ*_del_, which is at least |*q*| · *Δ*_match_ + |*Seeds*| · *δ_min_* by plugging in ndel from Eq. (3). This is an upper bound for *h*〈*v*, *i*〉, which we observe after maximizing *h*〈*v*, *i*〉 by substituting *i* = 1 and *misses* = |*Seeds*| into Eq. (1), which concludes the proof in this case.

Otherwise, *A* contains less than *n_del_* deletions. If we interpret *A* as a path in 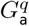, we first observe that *A* must spell *q*[*i*:]. Thus, *A* must in particular also contain all seeds *S_j_* ∈ *Seeds_≥i_* as substrings. We then split A into *subalignments A*__1_, *A*_0_,…, *A_p_*, selected such that *A*_0_,…, *A*_*p*–1_ spell the seeds *s_j_* ∈ *Seeds_≥i_*, and *A*__1_ and *A_p_* spell the prefix and suffix of *q*[*i*:] which do not cover any full seed.

This ensures that we can compute a lower bound on the cost of A as follows:

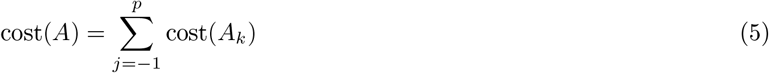

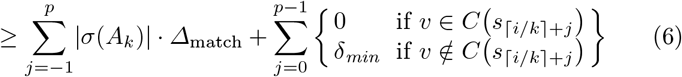

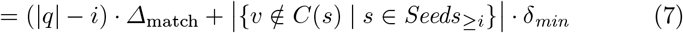

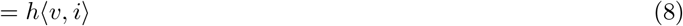

Here, Eq. (5) follows from our decomposition of A. If we ignore the right-hand side in Eq. (6) (right of “+”), the inequality follows because matching all letters is the cheapest method to align any string. The right-hand side follows from a more precise analysis for subalignments *A_k_* that spell a seed *s*_⌈*i*/*k*⌉+*j*_ without a corresponding crumb in *v*. The absence of such a crumb indicates that no exact match of *s*_⌈*i*/*k*⌉+*j*_ in *G*_r_ can be reached within less than *i* + *n_del_* steps from *v*. However, because *A* contains less than *n_del_* deletions, *A_k_* must start within less than *i* + *n_del_* steps from *v*. Thus, *A_k_* does not align *s*_⌈*i*/*k*⌉+*j*_ exactly, meaning that it introduces a cost of at least *δ_min_*.

Eq. (7) follows from observing that *A*__1_,…, *A_p_* have a total length of |*q*| — *i*, and observing that the right-hand sum adds up *δ_min_* for every expected but missing crumb. Finally, Eq. (8) follows from our definition of *h*〈*v*, *i*〉, concluding the proof.

#### Lemma 1 (Admissibility when skipping crumbs)

*The seed heuristic remains admissible when crumbs are skipped in the trie.*

*Proof.* Consider a reference graph with a match of seed s starting in node w. Now, consider a node *v* that cannot reach *w* using more than *i* – *D* – *n_ins_* edges. We can then show that a trie node *v*′ with a path to *v* does not require a crumb for the match of *s* in node *w*.

Specifically, any path from *root* through nodes *v*′ and *v* to node *w* has total length greater or equal *i* – *n*_ins_. Thus, matching *s* at *w* requires at least *n*_ins_ insertions. Hence, the cost of such a path is at least *n*_ins_ · *Δ*_ins_ = |*q*| · *Δ*_match_ + *n* · *δ_min_*. Observing that this is an upper bound for *h*〈*v*, *i*〉 concludes the proof.

### A.3 Versions, commands, parameters for running all evaluated approaches

In the following, we provide details on how we executed the newest versions of the tools discussed in §4:

#### Executing AStarix

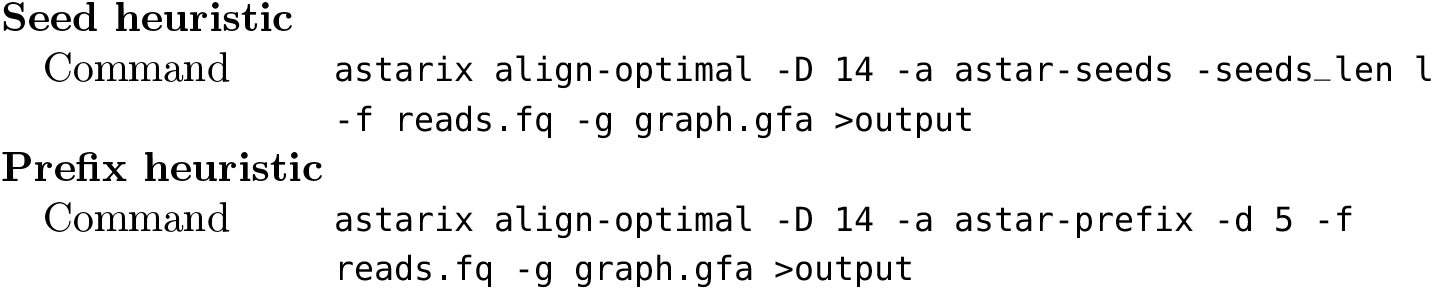

#### Executing other tools

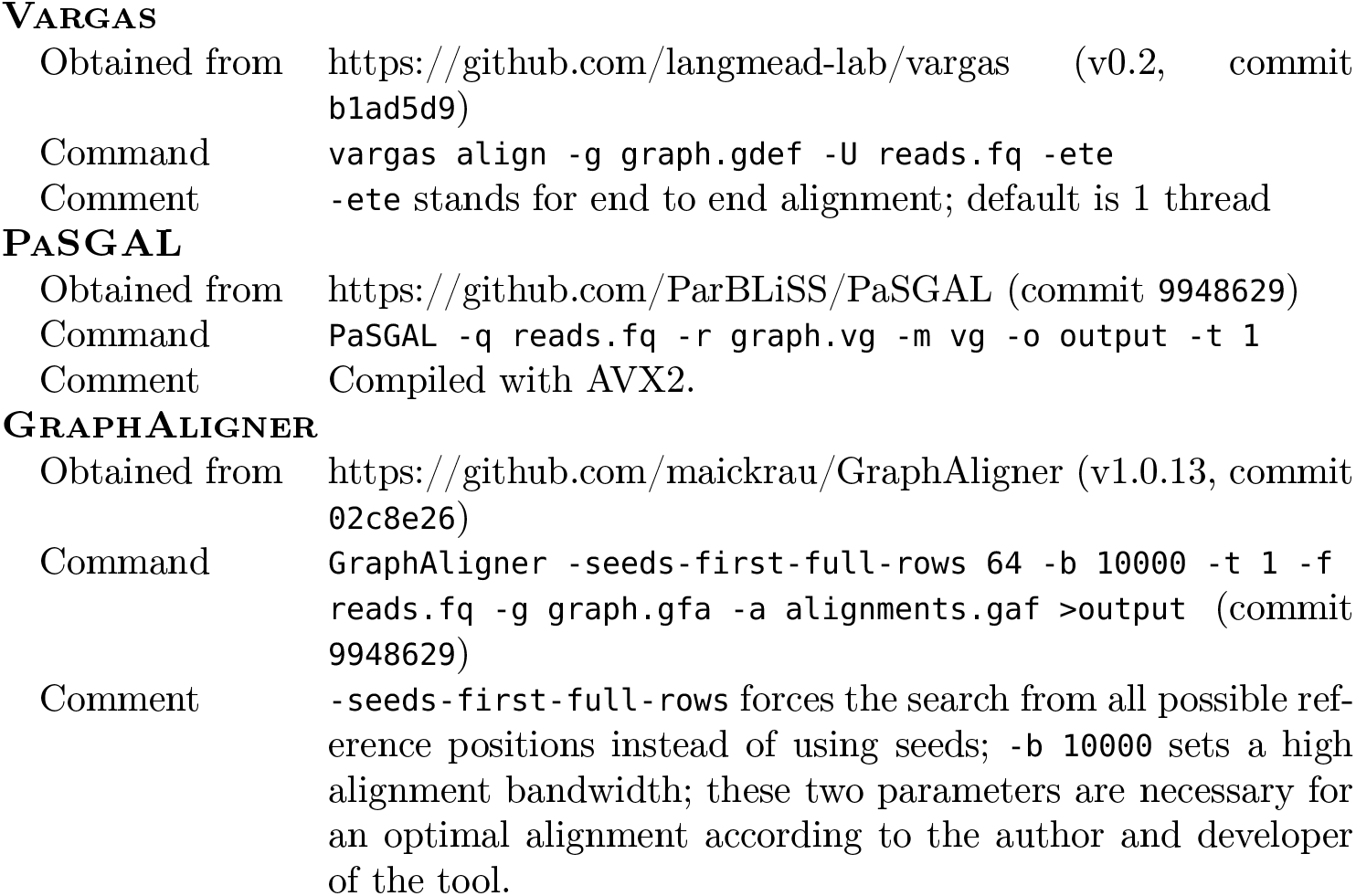

#### Simulating reads

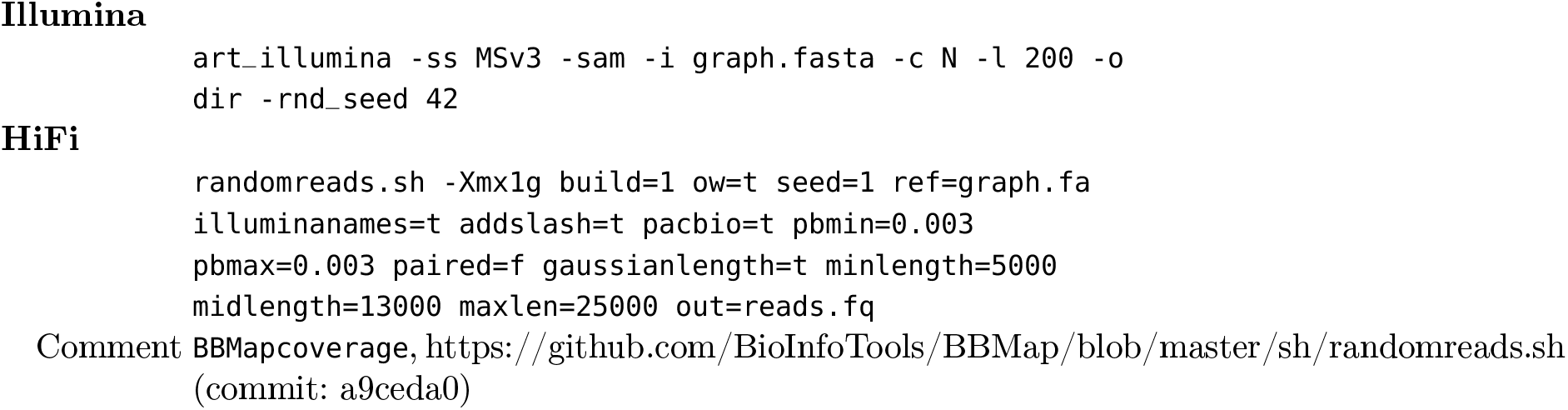

### A.4 Notations

Table 2 summarizes the notational conventions used in this work.

**Table 2.**
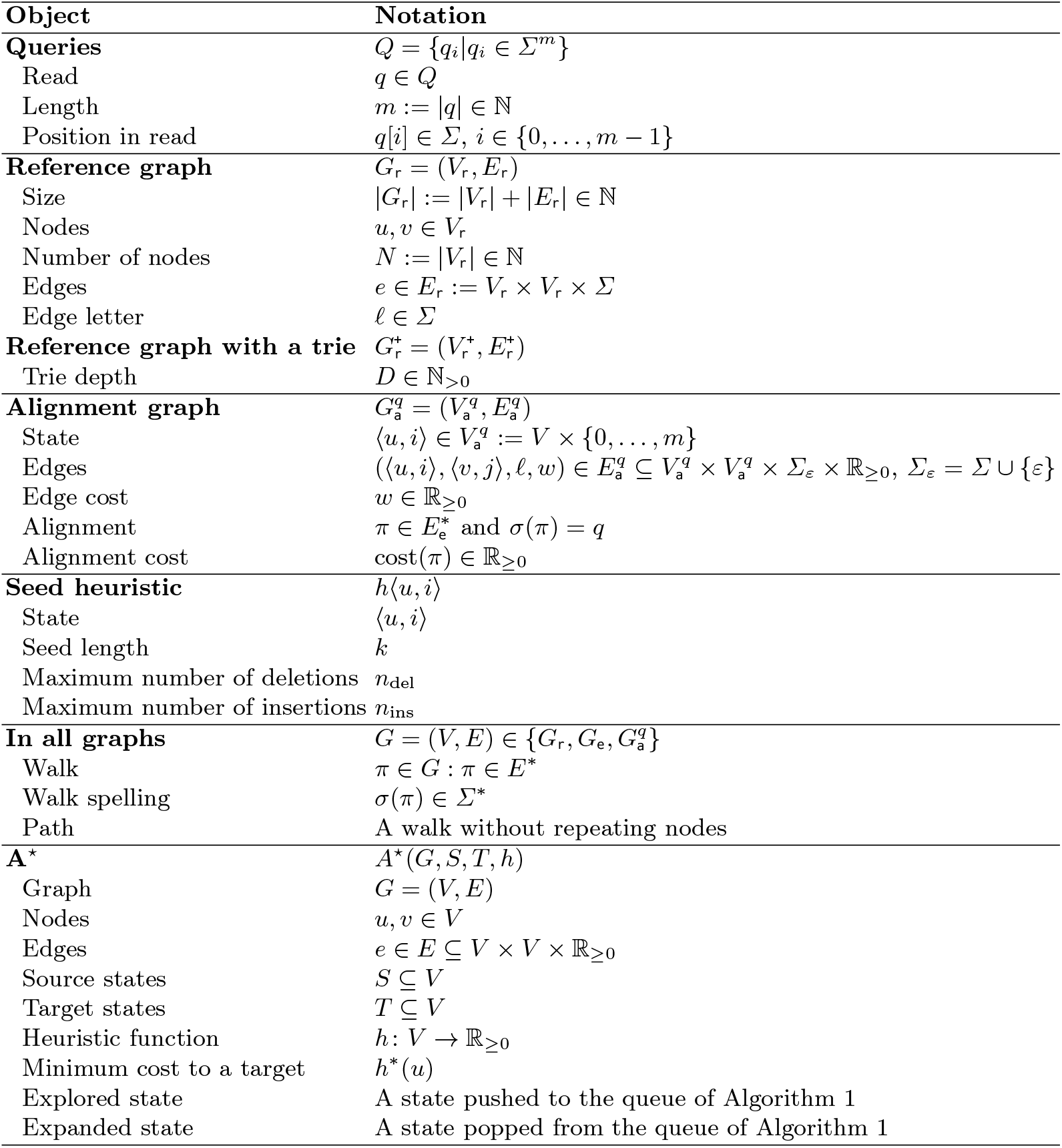
Notational conventions.

## Footnotes

1 Depending on how the A^⋆^ algorithm handles tie-braking, different sets of states could be explored. For simplicity, we show all states that *could potentially* be explored.

2 https://github.com/BioInfoTools/BBMap/blob/master/sh/randomreads.sh

3 https://snakemake.readthedocs.io/en/stable/

